# Disentangling Age and Schooling Effects on Inhibitory Control Development: An fNIRS Investigation

**DOI:** 10.1101/2021.07.06.451315

**Authors:** Courtney McKay, Sobanawartiny Wijeakumar, Eva Rafetseder, Yee Lee Shing

**Affiliations:** Division of Psychology, University of Stirling, Scotland; School of Psychology, University of Nottingham, UK; Department of Psychology, Goethe University Frankfurt, Germany; Center for Individual Development and Adaptive Education of Children at Risk (IDeA), Frankfurt, Germany; Department of Psychology, University of Konstanz

**Author notes:** Correspondence should be addressed to Yee Lee Shing, Department of Psychology, Goethe University Frankfurt, Frankfurt am Main, Germany.

## Abstract

Children show marked improvements in executive functioning (EF) between 4 and 7 years of age. In many societies, this time period coincides with the start of formal school education, in which children are required to follow rules in a structured environment, drawing heavily on EF processes such as inhibitory control. This study aimed to investigate the longitudinal development of two aspects of inhibitory control, namely response inhibition and response monitoring and their neural correlates. Specifically, we examined how their longitudinal development may differ by schooling experience, and their potential significance in predicting academic outcomes. Longitudinal data was collected in two groups of children at their homes. At T1, all children were roughly 4.5 years of age and neither group had attended formal schooling. One year later at T2, one group (P1, n = 40) had completed one full year of schooling while the other group (KG, n = 40) had stayed in kindergarten. Behavioural and brain activation data (measured with functional near-infrared spectroscopy, fNIRS) in response to a Go/No-Go task and measures of academic achievement were collected. We found that P1 children, compared to KG children, showed a greater change over time in activation related to response monitoring in the bilateral frontal cortex. The change in left frontal activation difference showed a small positive association with mathematical ability, suggesting certain functional relevance of response monitoring for academic performance. Overall, the school environment is important in shaping the development of the neural network underlying monitoring of one own’s performance.

**Research Highlights:**

- Using a school cut-off design, we collected longitudinal home assessments of two aspects of inhibitory control, namely response inhibition and response monitoring, and their neural correlates.
- For response monitoring, P1 children showed a greater difference over time in activation between correct and incorrect responses in the bilateral frontal cortex.
- The left frontal activation difference in P1 children showed a small association with mathematical ability, suggesting some functional relevance of response monitoring for academic performance.
- The school environment plays an important role in shaping the development of the neural network underlying monitoring of one own’s performance.

---

The developmental period of transitioning from kindergarten to formal schooling is characterized by remarkable improvements in cognitive functions. As children prepare for and settle into school and classroom environments, they are increasingly expected to orchestrate and exert control over their own thoughts and behaviors, in accordance to goals and context – a set of skills collectively known as executive functioning (EF; Diamond, 2013). In this study, we investigated the longitudinal development of a key component of EF, namely inhibitory control and its neural correlates, how these differ by schooling experience, and their potential significance in predicting academic outcomes.

There is accumulating evidence to suggest inhibitory control, the capacity to interrupt a prepotent response and enact an alternative less salient response associated with goal attainment, may play a key role in determining school readiness (Müller et al., 2008) as well as predicting future academic achievement (Blair & Razza, 2007a; Duckworth et al., 2019; Gawrilow et al., 2014; McClelland et al., 2014; Smith-Donald et al., 2007; Son et al., 2019). For instance, Bierman and colleauges (2008) found that, in a sample of typically developing preschool children, tasks of working memory and inhibitory control predicted emerging literacy skills. This finding is an agreement with Blair and Razza (2007), who examined the role of self-regulation in relation to emerging academic abilities in 3- to 5-year-old children. While several aspects of self-regulation predicted certain academic outcomes, inhibitory control made independent contributions to all three measures of academic ability (mathematical knowledge, letter knowledge, and phonemic awareness). The authors suggested that the ability to inhibit distracting or irrelevant information while reading or when faced with a numerical problem may be a contributing factor to success, over and above specific knowledge of problem solutions. For example, inhibitory control may allow children to consider multiple dimensions of a problem, rather than focusing on the most salient or recent aspects.

While inhibitory control prior to starting school may play an important role in predicting future academic success, the school environment itself may play an equally important role in shaping these skills. In school, children are required to follow classroom rules, sit still, and pay attention for a large portion of the lessons while suppressing any distractions that may interfere with their learning (Bierman et al., 2008). These demands draw heavily on inhibitory processes. Therefore, it is conceivable that the environment of formal schooling may advance the development of inhibitory control, in comparison to kindergartners that tend to be more play-oriented (Morrison et al., 1997).

To estimate the causal effects of schooling on cognitive development is not trivial, as schooling and development are confounded in time. The cut-off design (for a review, see Morrison et al., 2019) is an effective longitudinal method for examining unique schooling effects by taking advantage of arbitrary school cut-off dates. This method compares children who are similar in age, but due to fixed entry dates, are enrolled into different school years. Previous studies with a cut-off design found causal, beneficial effects of schooling on aspects of literacy (Morrison et al., 1995; Varnhagen et al., 1994) and numeracy (Bisanz et al., 1995; Christian et al., 2000). Recent years have seen a growth in research examining schooling-related effects on more basic cognitive processes, such as EF, given the associations shown between its subcomponents with academic achievement (Morrison et al., 2019). However, the findings here are mixed. For instance, Burrage et al. (2008) assessed inhibition in two groups of 5-year-old children born within 4 months of each other during the fall and spring semesters of the school year. The researchers found no significant difference in performance between children who had attended school and those who had stayed in kindergarten. On the other hand, Kim and colleauges (2021) used a school cut-off design to examine performance on an inhibitory control task in 4- to 7-year-old children. There was a significant difference between first grade children and kindergarten children, with kindergarteners showing greater improvements across the year. However, this result should be interpreted with caution for several reasons. First, initial differences existed between the two groups at the start of the year, with the first graders outperforming the kindergarteners at baseline. Hence, it is unclear whether the kindergarteners improved more from the experience of kindergarten or were just catching up in performance. Second, the difference between first grade and kindergarten children across time was no longer significant, when comparing children born within one month of the cut-off date as opposed to two months.

Despite the growing interest in how schooling may influence various aspects of basic cognition, there have been very few neurodevelopmental investigations. The only longitudinal inquiry into schooling-effects on neural correlates of inhibitory control was conducted by Brod and colleauges (2017). Using a cut-off design, fMRI data was collected on 5- and 6-year-old children while they completed a go/no-go task. This study sought to uncover schooling-related effects in *response inhibition*, and thus focused on activation for successfully inhibited (no-go) and successfully executed (go) trials. While no group differences in activation were found during correct no-go trials, a larger increase in activation in the right superior posterior parietal cortex (PPC), an area associated with sustained attention, was found for correct go trials, only in children who attended school. The authors concluded the increased engagement of the PPC may reflect a direct effect of the schooling experience, where children are required to pay attention for extended periods of time in classrooms.

Although trials with correct responses have traditionally been the focus of analyses in a go/no-go task, a separate literature have highlighted a unique pattern of activation in response to errors. First recognized by ERP researchers (Falkenstein et al., 1991; Gehring et al., 1993), the negative and positive components that arise following an incorrect response to a no-go trial are referred to as the error-related negativity (ERN) and error-related positivity (Pe). These components presumably reflect a network of structures, including the anterior cingulate cortex (ACC) and lateral prefrontal cortex (LPFC), and are thought to reflect error detection and/ or conflict resolution processes associated with *response monitoring* (Grammer et al., 2014; Kim et al., 2016).

Interestingly, response monitoring is one of the components of cognitive control that has been linked to academic success (Denervaud, Knebel, et al., 2020; Kim et al., 2016), and its deficits are associated with developmental disorders including ADHD (Groom et al., 2013). To be successful in school, children must monitor their own progress, detect errors when they occur, and subsequently adapt their own behaviour. In comparison to kindergarten, teachers in school classrooms also provide more directive feedback on the accuracy of children’s schoolwork, possibly shaping their sensitivity to errors (Denervaud, Knebel, et al., 2020). Relating response monitoring and schooling, Grammer et al. (2014) administered a go/no-go task to a sample of 3- to 7-year-old children and found that Pe was sensitive to age-related change during the school transition period, where older children exhibited a larger Pe than younger children. Further, Kim et al. (2016) administered a go/no-go task alongside two measures of academic achievement; math and reading. Using a multiple regression analysis, they found that stronger reading and math skills predicted a larger Pe but did not predict the ERN. Thus, the authors concluded that the Pe, rather than the ERN, may be associated with academic achievement. Most developmental research in response monitoring has been conducted using EEG, with a handful of studies that have used fMRI (Denervaud, Fornari, et al., 2020; Rubia et al., 2007). Specifically, Rubia and colleauges (2007) compared brain activation between adults and children while they completed a modified stop task. During unsuccessful stop trials (contrasted with successful go trials), adults and children showed similar activation in the medial prefrontal cortex, anterior, and posterior cingulate gyrus. However, adults showed increased activation compared to children in the ACC. Thus, converging evidence from fMRI and EEG investigations has identified neural signatures of response monitoring after committing error, and highlights the involvement of a network of frontal regions.

Based on the review above, there are several questions remained that the current study aimed to address. First, although Brod and colleauges (2017) reported that one year of formal schooling results in increased engagement of the PPC, it is unknown whether this increase predicts academic achievement. Previous studies that have investigated the link between response inhibition and academic achievement have been strictly correlational. Thus, any causal links between the two remain to be demonstrated. Second, it is unknown whether entering formal education causally impacts the frontal networks underlying response monitoring. None of the studies that examined response monitoring and schooling utilized a cut-off design. To fill in these knowledge gaps, we conducted a study in Scotland with a modified cut-off design. Rather than comparing children born several months before vs. after a cut-off date, all children in the current study were born in January and February of one year. This was possible because in Scotland, school commencement dates fall in August, with the school-starting cohort consisting of children born between the beginning of March in one year (aged 5.5) and the end of February (aged 4.5) of the following year. However, parents of children born in January and February can choose to enroll their child into school or defer their entry until the following year, and these requests are automatically approved. Thus, the current study compared two groups of children across time: one group enrolled into school as soon as they were eligible and completed one year of primary school (P1), and the other group deferred their school entry and stayed in kindergarten (KG). At timepoint 1 (T1) children in both groups were 4.5-years-old and in kindergarten. At timepoint 2 (T2), children in both groups were 5.5-years-old, but P1 children had completed one full year of schooling while KG children had completed another year of kindergarten.

Our first question sought to determine whether entering formal schooling leads to increased engagement of the neural networks underlying response inhibition and response monitoring. To answer this question, we employed a portable functional near-infrared spectroscopy (fNIRS) system, which allowed us to collect data on children in their homes (see more details in McKay et al., 2021). This system has several advantages over other imaging modalities as it is non-invasive, cost-effective, portable, and fairly easy to use with young children. Our second question inquired whether schooling-specific improvements in response inhibition and/ or monitoring, if any, would be associated with improvements in academic achievement^1^. In line with findings by Brod et al. (2017), we predicted both groups would show improvements in response inhibition, with P1 children showing a larger increase in parietal activation associated with sustained attention as a result of schooling. Further, based on research suggesting a link between response inhibition and future academic success (Blair & Razza, 2007; Gawrilow et al., 2014; McClelland et al., 2014; Smith-Donald et al., 2007; Son et al., 2019), we predicted the schooling-specific increase in parietal activation in the P1 children would be associated with larger improvements in academic achievement. Next, we predicted P1 children would, over time, show stronger response monitoring after committing error (i.e., acting wrongly based on prepotent response), and thus show a stronger change in activation in the frontal cortex in response to error trials. Lastly, based on work by Grammer et al. (2014), we predicted schooling-specific changes in response monitoring in the P1 children would be associated with improvements in academic achievement.

## Method

### Participants

Participants were recruited through gateway organizations such as nurseries and leisure centers. Parents of eligible children contacted the research team to schedule a testing session. All children had normal or corrected to normal vision, no history of colour-blindness, no neurological conditions, and were born full term (>37 weeks) with an uncomplicated birth. Parents and children provided informed consent prior to testing. The research was approved by the General University Ethics Panel (GUEP 375a) at the University of Stirling.

Children were tested in their home on two separate occasions, across two consecutive years. At T1, 95 4.5-year-olds were recruited for the study. Fifteen children were excluded from all analyses; 11 children interfered with the fNIRS set-up (pulled the cap off) before the completion of the task, four provided unusable data (two children had thick hair that led to poor signal quality, and data from two children was lost due to experimenter error). Hence, a total of 80 children (39 females, *Mage* at T1 = 53.5 months, *SD* = 1.2, range = 5 months) provided potentially usable fNIRS data at T1 (see further analysis-specific criteria below). All 80 children agreed to take part at T2 (39 females, *Mage* at T2 = 65.5 months, *SD* = 1.2, range = 5 months). Of these children, 40 (24 females, *M*age at T2 = 65.6 months, *SD* = 1.1, *range* = 5 months) attended P1 in between the two timepoints, and 40 (15 females, *M*age at T2 = 65.4 months, *SD* = 1, *range* = 4 months) remained in KG.

### fNIRS analysis exclusion (see Figure 1)

#### Response Inhibition

Two children were excluded from the response inhibition fNIRS analyses for contributing fewer than six usable correct no-go trials across both timepoints and five children were excluded for providing incomplete data (two children refused to complete the task and data from three children was corrupted). Hence, a total of 73 children contributed longitudinal data for the response inhibition fNIRS analyses. Of these children, 37 were in P1 group and 36 were in KG group.

#### Response Monitoring

Fifteen children were excluded from the response monitoring fNIRS analyses for contributing fewer than six usable incorrect no-go trials across both timepoints, and four children were excluded for providing incomplete data (one child refused to complete the task and data from three children was corrupted). A total of 61 children contributed to the response monitoring fNIRS analyses. Of these children, 30 were in P1 group and 31 were in KG group.

### Behavioural exclusion

#### Vocabulary

Three children were excluded from the vocabulary analyses (two refused to do the task and data from one child was lost due to experimenter error). 77 children contributed to the final vocabulary analyses. Of these children, 39 were in P1 group and 38 were in KG group.

#### Numeracy

Six children were excluded from the numeracy analyses (five refused to complete the task and data from one child was lost due to experimenter error). 74 children contributed to the final numeracy analyses. Of these children, 37 were in P1 group and 37 were in KG group.

#### School achievement packs (T2 only)

No children were excluded on either the math or phoneme pack.

### Experimental Task

#### Cats-and-Dogs Task

The cats-and-dogs task (CDT), adapted from Brod et al. (2017), was used to measure response inhibition and response monitoring in children – see Fig 2. The task was run in E-prime V.3 software on a HP laptop with a 14-inch screen. During “go” trials, children saw a picture of a dog and were supposed to press a button (spacebar). During “no-go” trials, children saw a picture of a cat and were supposed to withhold pressing a button. To ensure children understood the rules, the session began with 3 blocks of practice that progressively allowed less time to response. During the practice, children were reminded of the rules if they made a mistake. Performance on the practice runs was not included in final analyses. After children completed all practice blocks, the test session, consisting of two runs began. The first run was comprised of 59 trials: 44 go trials and 15 no-go trials. Run 2 was compromised of 69 trials: 52 go trials and 17 no-go trials. Pictures of cats and dogs were presented for 500ms, followed by a fixation cross as jitter that ranged in duration from 2 to 8 seconds. Responses made during stimuli presentation or during the fixation cross period were recorded. The order of presentation of go and no-go trials was pseudorandom, with the constraint that no-go trials were preceded equally often by 1, 2, 4 or 5 go trials.

**Fig. 1.**
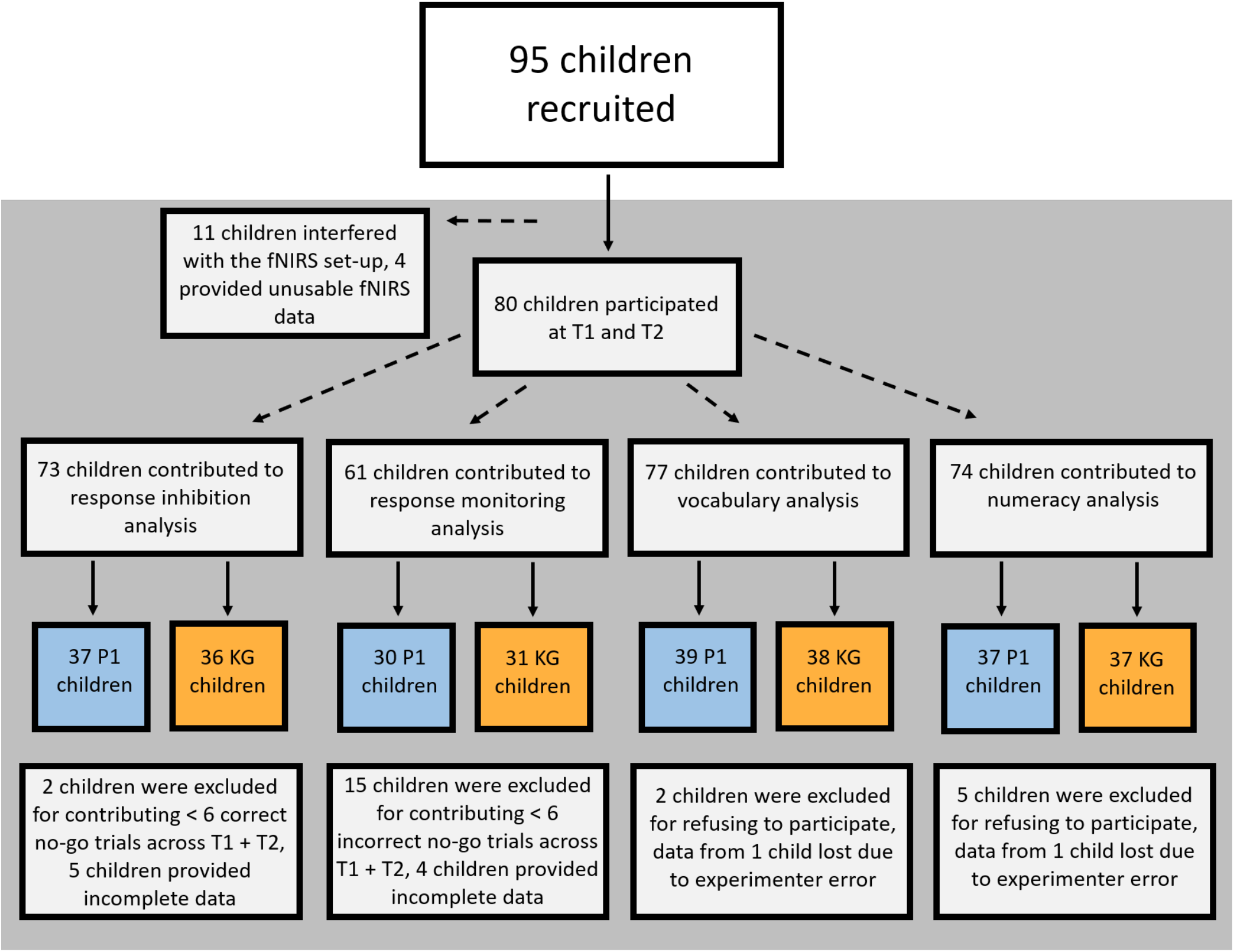
Schematic figure for participant recruitment and data exclusion.

**Fig. 2.**
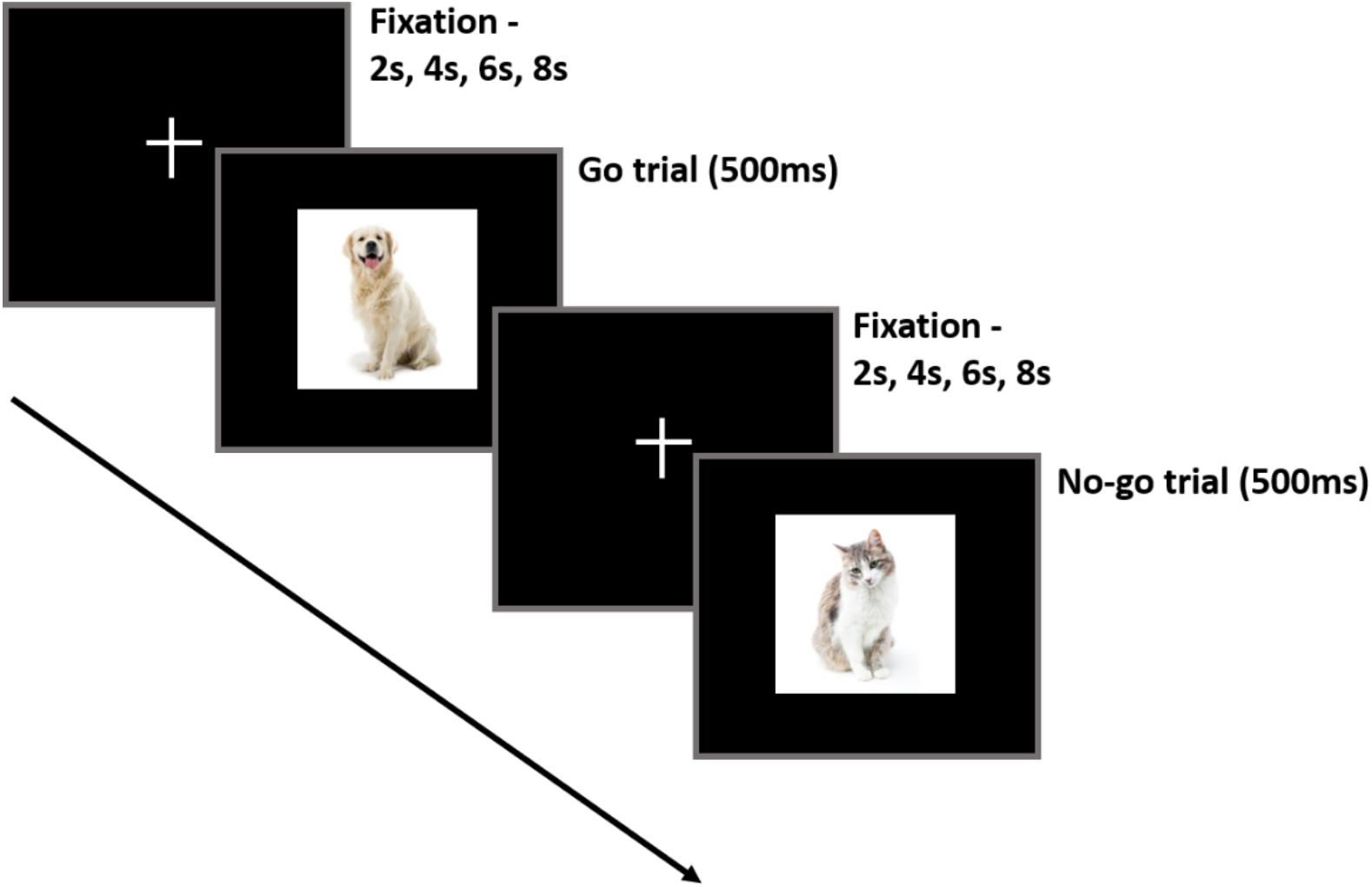
Trial structure of the Cats-and-Dogs Task (CDT).

### Academic Performance Measures

#### Vocabulary Task

The vocabulary subset of the Wechsler Preschool and Primary Scale of Intelligence (Warschausky & Raiford, 2018) was used to assess word knowledge. The task included 3 picture items and 20 verbal items. During the picture items, children were presented with 3 consecutive pictures of objects (car, scissors, banana) and asked to name each object. If a child incorrectly named the first object (car), they were corrected. Feedback was not provided for the other two picture items. For the verbal items, children were required to provide verbal definitions of words. Corrective feedback was given for the first two verbal items if a child did not receive a perfect score. No feedback was provided for the remaining verbal items. In accordance with the manual, if a child’s response was unclear or too vague, the experimenter prompted the child by asking, “What do you mean”, or “Tell me more about it”, or some other neutral query. The test was discontinued if a child gave three consecutive incorrect responses. The picture and verbal items were summed to provide a total vocabulary score (out of 43) at each timepoint.

#### Numeracy Task

The numeracy screener developed by Nosworthy and colleagues (2013) was used to assess basic numeracy skills. Children were required to compare pairs of magnitudes ranging from one to nine and judge which was larger. Magnitudes were represented symbolically (56 digit pairs) and non-symbolically (56 pairs of dot arrays). In both the symbolic and non-symbolic conditions, numerical magnitude was counterbalanced for the side of presentation. Dot stimuli were also controlled for area and density. Easier items were presented first, followed by more difficult items. Children were given one minute to complete each condition. The order of the two conditions were counterbalanced across participants. Children received one point for each correct answer. A final score was calculated at each timepoint by subtracting incorrect responses from correct responses.

### School achievement packs (T2 only)

Two measures of achievement were included to assess how much P1 children learned over the course of the first grade in terms of school content. The math pack contained 25 math questions, adapted from the Scottish Curriculum For Excellence teaching resources (*twinkl*, n.d.). The test was discontinued after 3 incorrect responses. The phonemes pack contained 20 questions assessing phonetic awareness, adapted from the Heggerty Phonemic Awareness Program (Heggerty, 2019). The pack included 10 items requiring the addition of a phoneme, and 10 items requiring the substitution of a phoneme. A final score for each pack was calculated by summing the correct responses.

### fNIRS data acquisition

fNIRS data were collected at 7.81 Hz using a NIRSport system 8×8 (8 sources 8 detectors) / release 2.01 with wavelengths of 850 and 760nm. Fiber optic cables carried light from the machine to a NIRS cap. Probe geometry was designed by collating regions of interest (ROI) from previous fNIRS and fMRI literature (Brod et al., 2017; Wijeakumar et al., 2015). Probe geometry consisted of four channels each on the left and right frontal cortices, and three channels each on the left and right parietal cortices (McKay et al., 2021). Note that short-source-detector channels were not used to regress scalp hemodynamics as all the channels were directed toward maximising coverage of the frontal and parietal cortices. Four cap sizes (50cm, 52cm, 54cm, and 56cm) were used to accommodate different head sizes. Source-detector separation was scaled according to cap size (50cm cap: 2.5cm; 52cm cap: 2.6cm; 54cm cap: 2.7cm and 56cm cap: 2.8cm). To synchronise behavioural and fNIRS data, a McDaq data acquisition device (www.mccdaq.com) was used to send information from the task presentation laptop to the fNIRS system.

### Procedure

Data was collected in each participant’s home. After arrival, the researcher measured the circumference of the child’s head and selected an appropriately sized fNIRS cap. Children were given an iPad to watch cartoons during the set-up. Once the cap was fitted to the child’s head, measurements were taken from the inion to the nasion and from the two peri-auricular points to make sure that the cap was centred. After the equipment was safely positioned, the instruction and practices for the CDT started, followed by the actual task. During the task, if children indicated that they made an error (e.g., pressing after a cat picture), the experimenter reassured the child and encouraged them to continue concentrating on the game. To keep children engaged, each test run contained different pictures of cats and dogs. To maintain motivation, children were also rewarded with a sticker after each run.

Once children completed the CDT, they were provided the iPad© to watch cartoons while the researchers removed the cap. After a short break, the testing proceeded with the vocabulary task, followed by the numeracy task. At T2, children were additionally tested on the phonemes pack and math pack. The order of the academic performance tasks were presented was counterbalanced across participants. Children were rewarded with stickers after completing each task, regardless of their performance. All children were remunerated with £10 and a toy upon completion of each time point measurement. As part of the overall procedure for the project, children also completed tasks on visual working memory, counterfactual reasoning, and associative memory, while parents filled in questionnaires collecting data on demographics, child behaviour, and life stress (data not included here, see details in McKay et al., 2021).

## Data analyses

### Behavioural analyses

Accuracy was calculated separately for each trial type (go and no-go) and test run (run 1 and run 2) and timepoint (T1 and T2). The following formula was used to calculate accuracy and RT at each timepoint to account for the different number of trials included in each run.

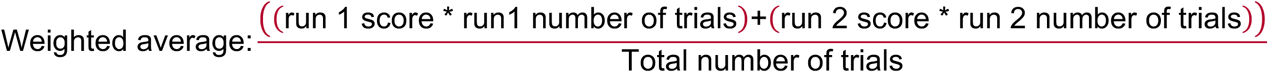

After computing the weighted averages, a corrected measure of accuracy (against response biases) was calculated for each subject, by subtracting no-go incorrect responses from go correct responses (Go_correct_-NoGo_incorrect_), separately at each timepoint.

### Outlier correction

All behavioural data were screened for outliers. To correct for longitudinal outliers, we used the Mahalanobis distance (MD) method. Further, we screened for outliers that were ±3 SDs from the mean at each timepoint. Three outliers were identified: two P1 children were removed from the phonemes pack analyses and one KG child was removed from the math pack analyses. No other outliers were identified.

### fNIRS preprocessing

fNIRS data were pre-processed using the Homer2 package (https://www.nitrc.org/projects/homer2/). Raw data were pruned using the *enPrunechannels* function (SNRthresh=2, SDrange=0.0 – 45). Signals were converted from intensity values to optical density (OD) units using the *Intensity2OD* function. Data was corrected for motion using the *hmrMotionCorrectPCArecurse* function, (tMotion=1, tMask=1, STDEVthresh=50, AMPthresh=0.5, nSV=0.97, maxlter=5, turnon=1). Data was scanned for motion artifacts using *hmrMotionArtifactByChannel* function (tMotion=1, tMask=1, STDEVthresh=50, AMPthresh=0.5). Then, the function *enStimRejection* (tRange=-1 to 3) was used to turn off stimulus triggers that contained motion artifacts. The data were band-pass filtered using *hmrBandpassFilt* to include frequencies between 0.016Hz and 0.5Hz. Using the function *hmrOD2Conc*, the OD units were converted to concentration units. To find trials that were outliers with respect to the average HRF, we used the function *hmrFindHrfOutlier* (tRange=-1 to 3, STDEVthresh=3, minNtrials=3). Lastly, the HRF was estimated using the ordinary least squares method with a modified gamma function with a square wave (*hmrDeconvHRF_DriftSS* function [tRange=-1 to 3, paramsBasis=0.1,0.5,0.5, rhoSD_ssThresh=0, flagSSmethod=0, driftOrder=3, flagMotionCorrect=0]).

### fNIRS group analyses

Oxygenated haemoglobin (HbO) and deoxygenated haemoglobin (HbR) beta values were extracted for each run (run 1 and run 2) and each condition (go cue correct trials, no-go cue incorrect trials, go cue incorrect trials, no-go cue incorrect trials, response go trials, response no-go trials). A weighted average was then calculated to account for the different number of trials included in each test run to produce one beta estimate per subject, per condition, per chromophore, and per timepoint.

### Response inhibition analyses

For the response inhibition analyses, we focused on HbO and HbR betas estimates for go cue correct trials and no-go cue correct trials. These beta values captured activation right after the onset of the stimulus. At T1, the mean number of correct trials included for P1s were 60 ± 4 go trials and 16 ± 1 no-go trials. The mean number of correct trials included for KGs were 66 ± 3 go trials and 18 ± 2 no-go trials. At T2, the mean number of correct trials included for P1s were 69 ± 4 go trials and 19 ± 1 no-go trials. The mean number of correct trials included for KGs were 75 ± 4 go trials and 18 ± 1 no-go trials.

### Response monitoring analyses

In the pre-registration, we initially only planned for analysis of response inhibition, focusing on correct responses on no go trials. However, based on consideration from the literature, we also investigated activation relating to response monitoring, namely contrasting erroneous responses on no-go trials against correct response on go trials. In both trial types a motor response was conducted, followed by no explicit feedback. Therefore, the post-processing of the erroneous response in the case of no-go trials is assumed to involve the detection of error and conflict, which should lead to more monitoring and careful responding in subsequent trials, consequently overall better performance on the task. Thus, for the response monitoring analyses, we focused on HbO and HbR betas estimates for response on go trials and response on no-go trials. These beta values captured activation at the onset of the child’s button press. At T1, the mean number of trials included for P1s were 61 ± 4 correct go trials and 12 ± 1 incorrect no-go trials. The mean number of trials included for KGs were 65 ± 4 correct go trials and 10 ± 1 incorrect no-go trials. At T2, the mean number of trials included for P1s were 69 ± 4 correct go trials and 9 ± 1 incorrect no-go trials. The mean number of trials included for KGs were 74 ± 4 correct go trials and 12 ± 1 incorrect no-go trials.

### Modelling framework

Univariate latent change score (LCS) models (Kievit et al., 2018; McArdle & Hamagami, 2004) were used to investigate the degree of change of the tasks with longitudinal data. All univariate models were set up as multi-group models, allowing the same model to be fitted for each group (P1 vs. KG) and later on parameter comparisons. Individual growth is captured by T1 (i.e., the intercept of *X1_T1* – Figure 3) and the latent change score factor (Δ*X1*), modelled as the difference between the initial observation and subsequent observation. Average group change across time is captured by the mean of the latent change score factor (*μ*_Δ*X1*_), and between-person differences in change are captured by the variance (*σ*^Δ*X1*^). Lastly, the covariance or regression parameter (*β*_*XT1*Δ*X1*_) determines to what extent the amount of change depends on scores at T1.

**Fig. 3.**
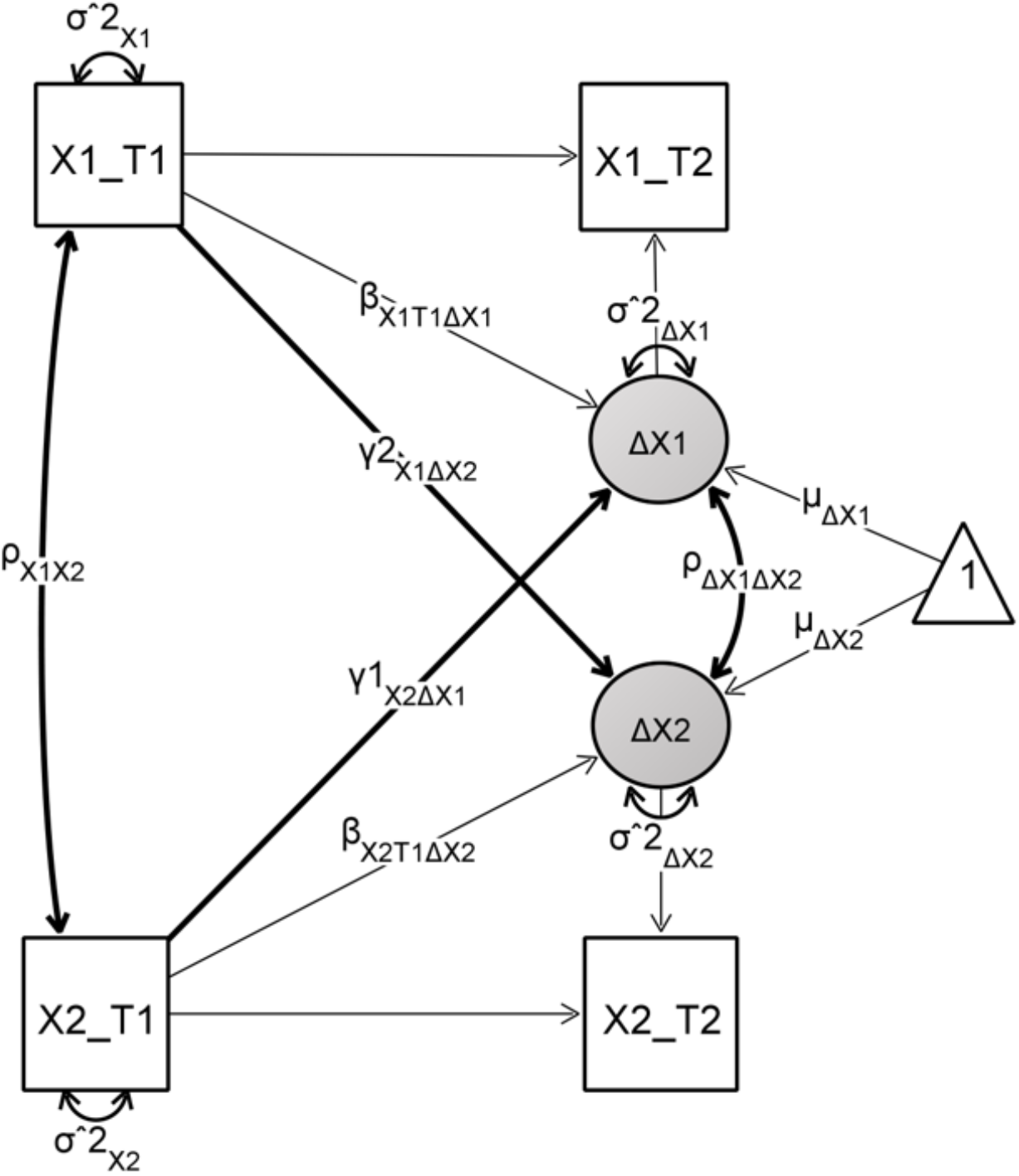
Graphical illustration of a bivariate latent change score model. Observed variables are depicted as squares and latent variables as circles. Variances are shown by two-headed arrows self, covariances are shown by two-headed arrows across variables, and regressions are shown by one-headed arrows. Figure created in Onyx (http://onyx.brandmaier.de).

With the inclusion of an extra domain, a univariate LCS model can be extended into a bivariate LCS model, allowing for testing of cross-domain coupling (see Figure 3). To determine whether scores at T1 in one domain (*X1*) are associated with scores at T1 in a second domain (*X2*), the intercept covariance (*ρ_X1X2_*) is estimated. To examine whether the change in *X1* is associated with the change in *X2*, the change covariance is estimated (*ρ*_Δ*X1*Δ*X2*_). Further, the coupling effect (*y2*_*X1*Δ*X2*_) determines whether the change in *X1* is a function of the starting point of *X2*, and vice versa (*y1*_*X2*Δ*X1*_). For the bivariate LCS model, as motivated by our second research question, only measures that showed schooling-specific effects, from response inhibition/monitoring on the one hand, and academic performance, on the other hand, were included.

### Model fit indices

Models were estimated in the lavaan software package in R (version 3.6.2, 2019; Rosseel, 2012). Full information maximum likelihood was used for model estimation. To test for significance of parameters of interest, equality constraint was made on the parameter and significance of change in model fit (compared to the just-identified free model) was assessed using the chi-square difference test (at p< .05). To account for any age and gender effects, these variables were added as covariates of no interest into all models.

## Results

### Behavioural results

#### Univariate LCS modelling

Four separate univariate models were fitted to each group (P1 and KG) with (1) corrected accuracy on CDT (*Go_correct_-NoGo_incorrect_*) (2) vocabulary scores (3) symbolic numeracy scores (4) non-symbolic numeracy scores. Raw mean performance levels are illustrated in Figure 4. Parameter estimates are shown in Table 1.

**Fig. 4.**
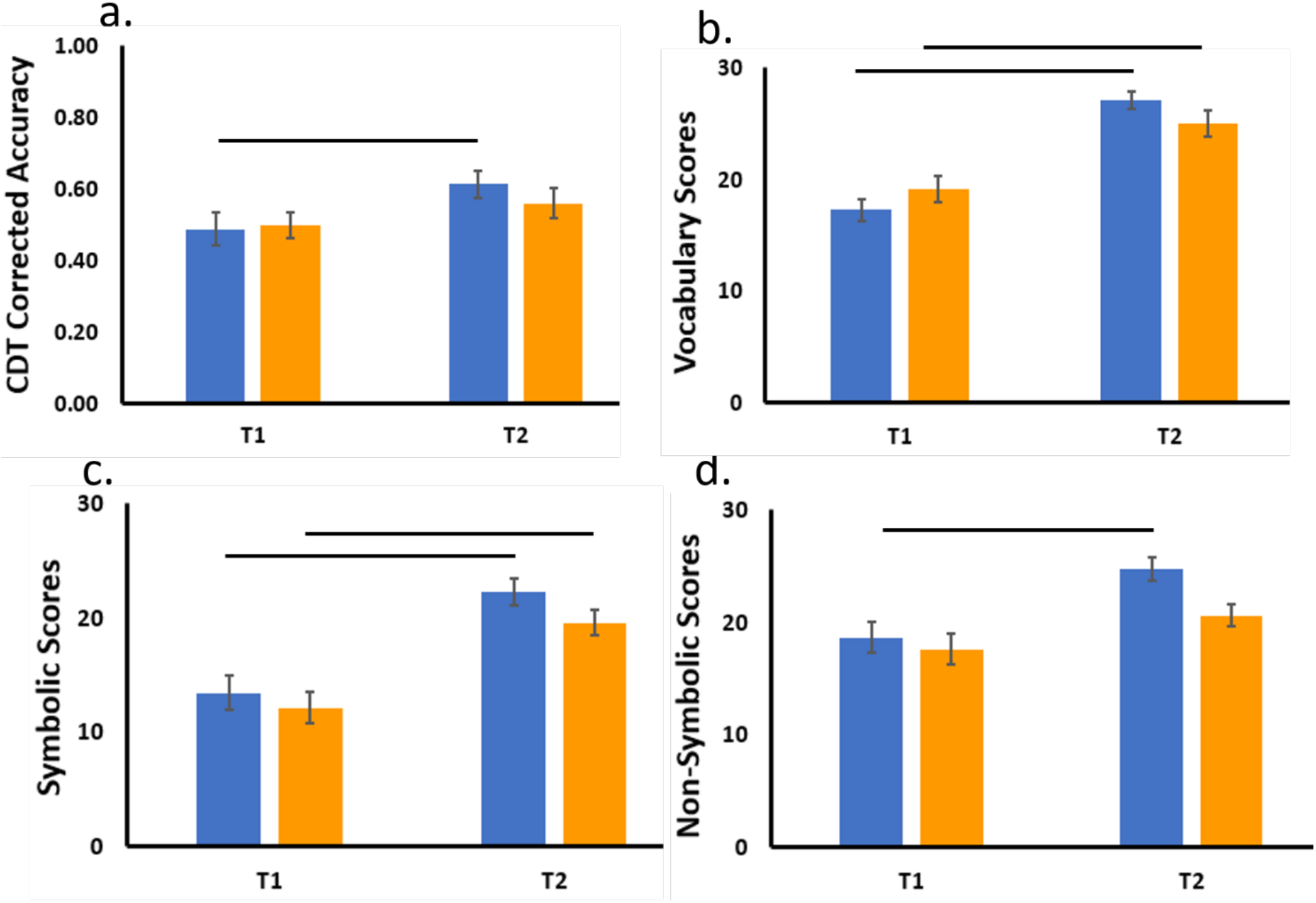
Behavioural estimates for the **(a)** CDT task (corrected accuracy based on Go_correct_-NoGo_incorrect_) **(b)** vocabulary task **(c)** numeracy task (symbolic) **(d)** numeracy task (non-symbolic). P1 children are shown in blue and KG children are shown in orange. denotes significance at *p* <.05 level (see text for the results of formal model comparison). Error bars show SEM.

**Table 1.**
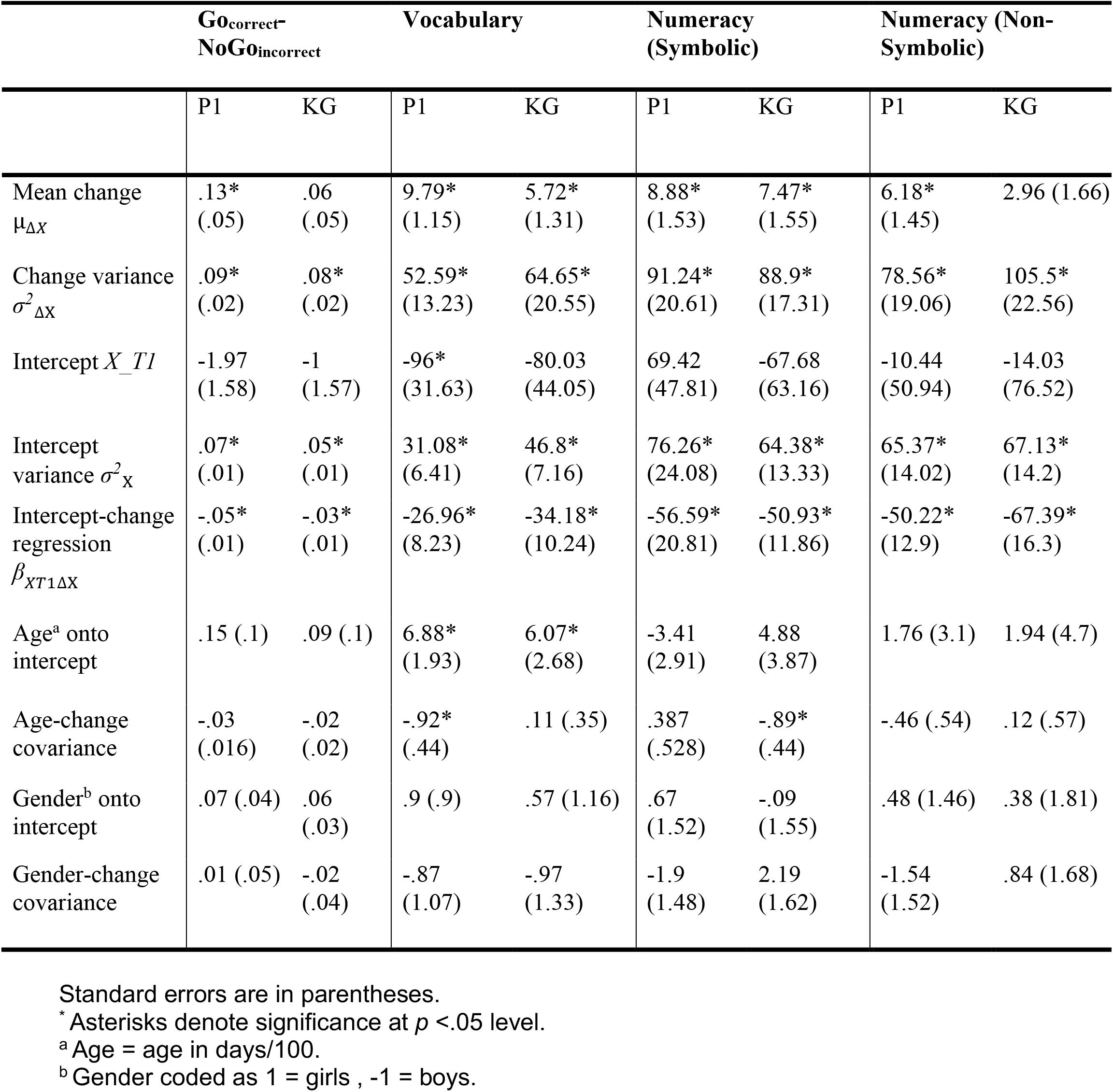
Parameter estimates for P1 children and KG children from four separate univariate models on the behavioral tasks.

##### CDT

P1 children showed a significant increase in corrected accuracy between T1 and T2, while KG children did not. However, when the change in corrected accuracy was constrained to be equal across groups, model fit was not significantly worse, Δx^2^ = 1.237, Δ*df* = 1, *p* = .266. There was also no significant worsening in model fit when the baseline scores at T1 were constrained to be equal across groups Δx^2^ = .189, Δ*df* = 1, *p* = .664. This suggests that P1 children and KG children started out with similar accuracy and changed comparably across the two timepoints, contrary to our hypothesis.

##### Vocabulary

Both P1 children and KG children showed a significant increase in vocabulary scores between T1 and T2. Constraining the change to be equal across groups led to a significant drop in model fit Δx^2^ = 5.001, Δ*df* = 1, *p* = .025, suggesting P1 children increased significantly more than KG children. No significant differences at T1 were found Δx^2^ = .084, Δ*df* = 1, *p* = .772. Therefore, P1 children and KG children started out with similar accuracy, but the improvement in P1 children on vocabulary knowledge was greater than the improvement in KG children.

##### Numeracy

For the symbolic condition, both P1 children and KG children showed a significant increase in scores between T1 and T2. No significant drop in model fit was found when the change was constrained to be equal across groups Δx^2^ = .413, Δ*df* = 1, *p* = .520. Further, no significant baseline difference was found when the scores at T1 were constrained to be equal across groups Δx^2^ = 3, Δ*df* = 1, *p* = .083. For the non-symbolic condition, P1 children significantly improved between the two timepoints while KG children did not. However, when the change was constrained to be equal across groups, no significant drop in model fit was observed Δx^2^ = 2.037, Δ*df* = 1, *p* = .154. Further, no significant drop in model fit was found after constraining T1 estimates to be equal across groups Δx^2^ = .002, Δ*df* = 1, *p* = .969. Thus, for both conditions of the task, P1 children and KG children started out with similar scores and they changed comparably between the two timepoints.

##### School achievement packs

Univariate models could not be fitted to the school achievement packs as they were only administered at T2, after P1 children had completed 1 year of schooling. Thus, simple t-tests were conducted to compare performance between P1 and KG children on these measures. As expected, we found that P1 children (Math: *M* = 30.1, *SD* = 6.6; Phonemes: *M* = 6.4, *SD* = 4) performed significantly better than KG children (Math: *M* = 23.9, *SD* = 6.5; Phonemes: *M* = 2.5, *SD* = 2.7) on both math and phonemes, respectively (t[77] = 4.233, *p* <.001; t[76] = 5.067, *p* <.001).

### fNIRS Results

fNIRS data were comprised of HbO and HbR beta values for each of the 14 channels. To reduce data dimension and focus subsequent analyses on effects that had a difference between HbO and HbR, an initial repeated measure ANOVA including chromophore (HbO, HbR) as a factor was run for each channel, using the Benjamini-Hochberg method to correct for multiple comparisons. For the response inhibition analyses, a repeated measures ANOVA with a within-subject factor of trial type (go correct, no-go correct) and chromophore (HbO, HbR) and a between-subjects factor of group (P1, KG) was run for each of the 14 channels. For the response monitoring analyses, a repeated measures ANOVA with a within-subject factor of trial type (go correct, no-go incorrect) and chromophore (HbO, HbR) and a between-subjects factor of group (P1, KG) was run for each of the 14 channels. We focused on significant interactions involving chromophore as a factor, and followed up with post-hoc analyses conducted on the HbO estimates.

#### Response inhibition analyses

Only channels that showed a significant interaction involving chromophore and that survived the Benjamini-Hochberg correction are reported. The interaction between trial type and chromophore was significant in channels overlying the right middle frontal gyrus (*F*[1,71] = 12.052, *p*=.001), the right inferior frontal gyrus (*F*[1,71] = 8.241, *p*=.005), the right supramarginal gyrus (*F*[1,70] = 7.932, *p*=.006), and the left supramarginal gyrus (*F*[1,71] = 11.876, *p*=.001). Following up on the interaction, post-hoc tests revealed that HbO activation for go correct trials was greater than activation for no-go correct trials (see Table 2). The remaining 3-and 4-way interactions between group, trial type, time, and chromophore were either not significant or did not survive the Benjamini-Hochberg correction.

**Table 2.** Response inhibition analysis: channels showing significant interactions between trial type (go correct and no-go correct) and chromophore. Significant post-hoc results are shown for HbO estimates.

| Channel No. | Brain areas (MNI labels) | Trial x Chromophore (HbO) |
| --- | --- | --- |
| Channel 1 | Right middle frontal gyrus | Go > No-go ($p=.007$) |
| Channel 2 | Right middle frontal gyrus |  |
| Channel 3 | Right inferior frontal gyrus | Go > No-go ($p=.006$) |
| Channel 4 | Right inferior frontal gyrus |  |
| Channel 5 | Left middle frontal gyrus |  |
| Channel 6 | Left middle frontal gyrus |  |
| Channel 7 | Left inferior frontal gyrus |  |
| Channel 8 | Left inferior frontal gyrus |  |
| Channel 9 | Right angular gyrus |  |
| Channel 10 | Right superior occipital gyrus |  |
| Channel 11 | Right supramarginal gyrus | Go > No-go ($p=.008$) |
| Channel 12 | Left inferior parietal lobule |  |
| Channel 13 | Left angular gyrus |  |
| Channel 14 | Left supramarginal gyrus | Go > No-go ($p=.004$) |

#### Response monitoring analyses

Only channels that showed a significant interaction with chromophore and that survived the Benjamini-Hochberg correction are reported. The interaction between trial type and chromophore was significant in channels overlying the right middle frontal gyrus (*F*[1,57] = 21.134, *p*<.001; *F*[1,57] = 15.341, *p*<.001), the right inferior frontal gyrus (*F*[1,57] = 19.023, *p*<.001), the left middle frontal gyrus (*F*[1,57] = 40.548, *p*<.001), the left inferior frontal gyrus (*F*[1,57] = 18.279, *p*<.001; *F*[1,57] = 10.769, *p*=.002), and the right supramarginal gyrus (*F*[1,56] = 6.773, *p*=.012). Following up on the interaction, post-hoc tests revealed that HbO activation for (erroneous) response at no-go trials was more negative than (correct) response at go trials (see Table 3).

**Table 3.** Response monitoring analysis: channels showing significant 2-way interaction between trial type (go correct vs. no-go incorrect) and chromophore, and 4-way interaction between group, trial type, time, and chromophore. Significant post-hoc results are shown for HbO estimates.

| Channel No. | Brain areas (MNI labels) | Trial x Chromophore (HbO) | Group x Trial x Time x Chromophore (HbO) |
| --- | --- | --- | --- |
| Channel 1 | Right middle frontal gyrus | Go > No-go, ($p < .001$) | P1 T1: Go > No-go, ($p = .048$)<br>P1 T2: Go > No-go, ($p = .001$)<br>KG T1: Go > No-go, ($p = .036$) |
| Channel 2 | Right middle frontal gyrus | Go > No-go, ($p < .001$) | P1 T1: Go > No-go, ($p = .013$)<br>P1 T2: Go > No-go, ($p < .001$) |
| Channel 3 | Right inferior frontal gyrus | Go > No-go, ($p < .001$) | P1 T1: Go > No-go, ($p = .004$)<br>P1 T2: Go > No-go, ($p < .001$)<br>KG T1: Go > No-go, ($p = .005$) |
| Channel 4 | Right inferior frontal gyrus |  |  |
| Channel 5 | Left middle frontal gyrus | Go > No-go, ($p < .001$) | P1 T1: Go > No-go, ($p = .001$)<br>P1 T2: Go > No-go, ($p = .002$) |
| Channel 6 | Left middle frontal gyrus |  |  |
| Channel 7 | Left inferior frontal gyrus | Go > No-go, ($p < .001$) | P1 T1: Go > No-go, ($p = .012$)<br>P1 T2: Go > No-go, ($p = .001$)<br>KG T1: Go > No-go, ($p < .001$) |
| Channel 8 | Left inferior frontal gyrus | Go > No-go, ($p = .025$) | |
| Channel 9 | Right angular gyrus |  |  |
| Channel 10 | Right superior occipital gyrus | | P1 T2: Go > No-go, ($p = .05$) |
| Channel 11 | Right supramarginal gyrus | Go > No-go, ($p = .042$) | |
| Channel 12 | Left inferior parietal lobule |  |  |
| Channel 13 | Left angular gyrus |  |  |
| Channel 14 | Left supramarginal gyrus |  |  |

A significant 4-way interaction between group, time, trial, and chromophore was observed in channels overlying the right middle frontal gyrus (*F*[1,57] = 10.198, *p*=.002; *F*[1,57] = 5.671, *p*=.021), the right inferior frontal gyrus (*F*[1,57] = 7.402, *p*=.009), the left middle frontal gyrus (*F*[1,57] = 9.912, *p*=.003), the left inferior frontal gyrus (*F*[1,57] = 5.897, *p*=.018), and the right superior occipital gyrus (*F*[1,56] = 5.976, *p*=.018). All post-hoc tests are shown in Table 3. Importantly, in the bilateral middle frontal gyrus and bilateral inferior frontal gyrus, P1 children showed greater negative activation for response at incorrect no-go trials than for correct go trials at both T1 and at T2. This was not the case for KG children, who only showed a difference in activation between these trials at T1. Therefore, the ANOVA revealed that the difference in activation between correct go trials and incorrect no-go trials across time differentiated P1 children from KG children. To relate these neural differences in response monitoring to behavior using the bivariate LCS models, an average difference in activation (go correct activation – no-go incorrect activation) was computed across channels of nearby regions that showed the significant 4-way interaction with similar patterns. Specifically, this led to two clusters covering the right frontal cortex (averaging channels 1, 2, and 3; see Figure 5a) and the left frontal cortex (averaging channels 5 and 7; see Figure 5b).

**Fig. 5.**
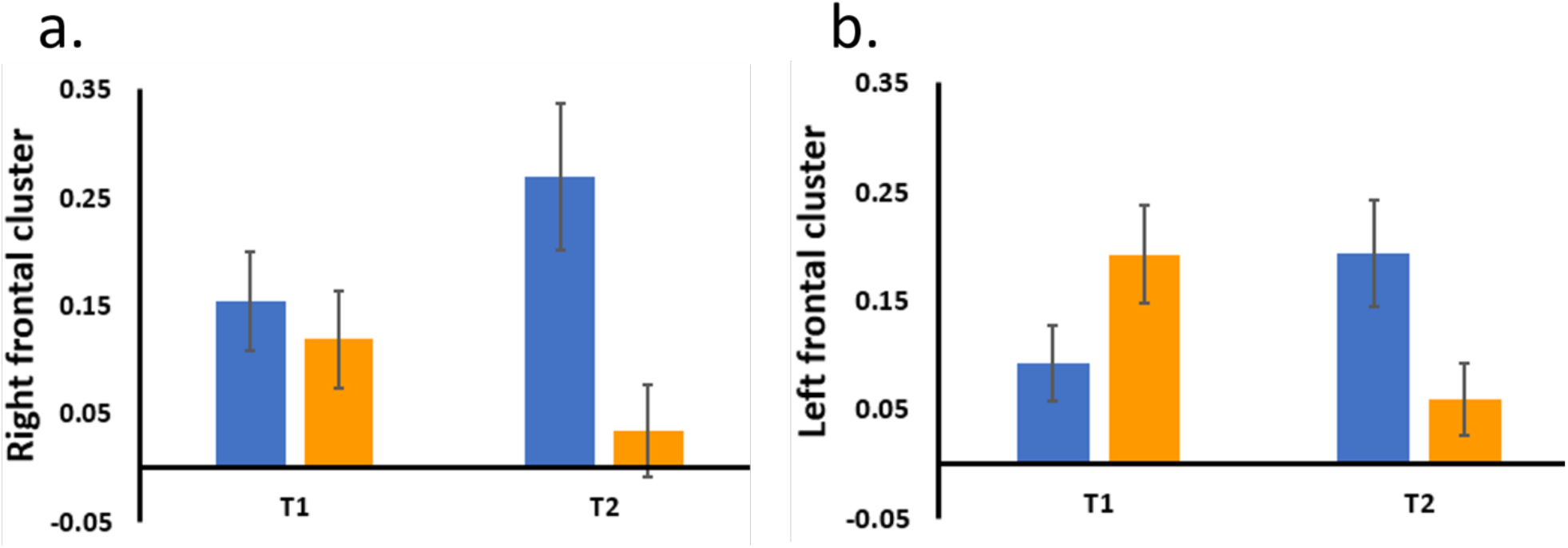
The difference in activation between go correct and no-go incorrect trials (response monitoring contrast) in the **(a)** right frontal cluster **(b)** left frontal cluster. P1 children are shown in blue and KG children are shown in orange. Error bars show SEM.

### Bivariate LCS modelling

As the first step, we tested the longitudinal coupling between activation difference in the two frontal clusters and corrected accuracy on the CDT task for both groups. This is mainly to verify the functional relevance of the two frontal clusters of response monitoring activation for overall task performance. Second, to address our second research question, we tested to what extent the response monitoring activations in the two frontal clusters, which showed diverging pattern of change in the two groups, could predict academic achievement. The longitudinal coupling between the activation difference with performance on the academic tasks was examined. Here, we focused on bivariate relationships of P1 children (since they were the only group that attended school). Bivariate relationships for KG children are shown in Supplementary Material.

#### Right frontal cluster and CDT corrected accuracy

Parameter estimates are shown in Table 4. For KG Children, corrected accuracy at T1 was positively correlated with the difference in activation in the right frontal cluster at T1. Namely, children who showed more difference in activation related to response monitoring had better performance. Constraining the baseline correlation at T1 to be 0 in KG children led to a significant drop in model fit, Δx^2^ = 10.707, Δ*df* = 1, *p* = .001. No other cross-domain parameters were significant.

**Table 4.** CDT bivariate couplings between (a) right frontal cluster and corrected accuracy (Go_correct_-NoGo_incorrect_)and (b) left frontal cluster and corrected accuracy (Go_correct_-NoGo_incorrect_), separately for P1 children and KG children.

|  | a. Right frontal cluster |  | b. Left frontal cluster |  |
| --- | --- | --- | --- | --- |
|  | P1 | KG | P1 | KG |
| Intercept covariance $\rho_{X1X2}$ | .65 (1.3) | 2.97* (.78) | 3.32* (1) | 1.83* (.82) |
| Right frontal onto corrected accuracy change $Y1_{X2\Delta X1}$ | 0 (0) | -.01 (0) | 0 (0) | 0 (0) |
| Corrected accuracy onto right frontal cluster change $Y2_{X1\Delta X2}$ | -53.5* (22.37) | 3.03 (20.73) | -10.1 (19.92) | 31.91* (14.6) |
| Change-change covariance $\rho_{\Delta X1\Delta X2}$ | .98 (1.41) | -.47 (1.11) | 1.35 (1.31) | 1.16 (.73) |

Standard errors are in parentheses.
\* Asterisks denote significance at $p < .05$ level.

For P1 children, corrected accuracy at T1 negatively predicted the change in the difference in activation in the right frontal cluster from T1 to T2. Thus, children with better performance at T1 showed less change in activation over time. However, constraining the coupling pathway to be 0 did not lead to a significant drop in model fit Δx^2^ = 3.776, Δ*df* = 1, *p* = .052. No other cross-domain parameters were significant.

#### Left frontal cluster and CDT corrected accuracy

For KG children, better corrected accuracy at T1 was correlated with higher difference in activation in the left frontal at T1. Constraining the baseline correlation at T1 to be 0 in KG children led to a significant drop in model fit, Δx^2^ = 5.028, Δ*df* = 1, *p* = .025. Furthermore, higher corrected accuracy at T1 predicted more change in the difference in activation in the left frontal. To follow up on this, the coupling pathway was constrained to be 0 in KG children, which led to a significant drop in model fit Δx^2^ = 4.492, Δ*df* = 1, *p* = .034.

For P1 children, similar to KG children, better corrected accuracy at T1 was correlated with higher difference in activation in the left frontal at T1. Constraining the baseline correlation at T1 to be 0 in P1 children lead to a significant drop in model fit, Δx^2^ = 5.536, Δ*df* = 1, *p* = .019. No other cross-domain pathways were significant. Taken together, in KG and P1 children, higher response monitoring activation difference in the left frontal cluster (additionally right frontal cluster for KG) was related to better overall performance in the inhibitory control task.

### Academic Achievement in P1 children

#### Right/left frontal cluster and academic achievement

Bivariate longitudinal models were fitted for the response monitoring activation in the right frontal (or left frontal, respectively) and (1) vocabulary scores (2) math pack and (3) phonemes pack. The longitudinal change in activation in the left frontal cluster was positively correlated with math pack scores at T2 (*p*=.04). To follow up on this finding, the coupling pathway was constrained to be 0, which led to a borderline significant drop in model fit Δx^2^ = 3.488, Δ*df* = 1, *p* = .062. No other cross-domain parameters were found to be significant in all other models.

## Discussion

The present study sought to examine to what extent one year of formal schooling shapes the development of neural processes underlying response inhibition and response monitoring, and establish whether these effects were related to academic achievement. First, we found that P1 children and KG children started out with similar corrected accuracy on the go/no-go task. Although P1 children, but not KG children, showed significant improvement on task accuracy over time, the magnitude of change between the two groups was statistically comparable. While we hypothesized that P1 children would show greater improvement than KG children across the year, our findings are in line with Brod et al. (2017) who also reported no group differences in response inhibition behaviour across the year. However, unlike Brod et al. (2017) and in contrary to our hypothesis, we also did not find any group difference in neural activation related to response inhibition (or parietal activation during go trials as in Brod et al. 2017).

Several methodological differences exist that may account for this inconsistency. First, children in the current study were between one to two years younger than the children in the Brod et al. (2017) study, due to national differences in school entry age. The first year of schooling may be set up to be less demanding and formally structured in countries where children start school at a younger age. Thus, the increase in parietal activation resulting from a schooling environment may only appear if there is a sufficiently large change in terms of demand and expectations transitioning from kindergartens to classrooms. Second, the current study and that of Brod et al. (2017) employed different modalities to record brain activation. The fNIRS channel-based analyses employed here may not have been as sensitive as the fMRI analyses conducted by Brod et al. (2017) to detect changes in activation in small clusters of voxels (as reported in that study). A potential way to improve upon this would be to conduct more targeted analyses. For instance, novel image reconstruction uses a head model to generate functional images of the fNIRS data, transforming surface level channel-based data into a volumetric representation within the brain (Forbes et al., 2021). This would allow for greater comparability with fMRI investigations. Another limitation of our research may be related to the longitudinal nature of the study. Longitudinal research with fNIRS carries the risk that the recorded areas do not remain consistent over time, particularly in development when children’s brains are actively developing and growing. While the risk may never be eliminated, steps were taken to limit the possibility that shifting head sizes would interfere with the recordings. Care was taken to ensure the appropriately sized cap was chosen for each child based on their head circumference. Furthermore, the experimenters took measurements as precise as possible to ensure the center of the cap was aligned with the center of the child’s head.

Second, for activation related to response monitoring, we found that P1 children, but not KG children, showed a greater difference after one year of schooling. As the response monitoring contrast was not part of the study pre-registration, it was important for us to first establish the functional relevance of the two frontal clusters (left and right middle/inferior frontal gyrus) that emerged from this contrast. Therefore, we tested the coupling between the difference in activation with performance on the CDT task, and we found that a greater response monitoring activation difference in the left frontal cluster was related to better performance in the inhibitory control task in both groups. For KG children, a similar relationship was also found for the right frontal cluster. This is in line with previous research reporting that a greater difference in activation between correct go and incorrect no-go trials reflects more efficient response monitoring (Grammer et al., 2014; Torpey et al., 2012), which may support better task performance. Previous adult fMRI studies have implicated a broader network of frontal regions subserving response monitoring. For example, Chevrier et al., (2007) administered a stop-signal task and found error-related activity in frontal regions including the right middle frontal gyrus and dorsal ACC. Furthermore, Edwards et al., (2012) administered a go/no-go task and combined ERP time courses and fMRI spatial maps allowing for the identification of brain regions that are associated with portions of the time course in the ERP data. They identified two components associated with significant activation in the bilateral middle frontal gyrus and caudal ACC, demonstrating that both regions are engaged during error processing. The authors argued the simultaneous involvement of both areas may reflect a post-error cognitive response, where conflict between the executed and supposedly correct response occurs via the caudal ACC and LPFC. Based on experimenter observations in the current study, this interpretation seems likely as children sometimes showed a reaction reflecting conflict after making an incorrect button press in a no-go trial. Children would either verbally indicated that they made a mistake (e.g., saying “oh no”) or show behavior of having committed an error (e.g., clasping hands over mouth, pulling hand away from keyboard).

Turning to schooling effect, In the two frontal clusters identified from the response monitoring contrast, P1 children showed a greater difference in activation across time than KG children. We posit that, across the first school year, P1 children show stronger response monitoring due to the nature of the schooling environment. In school, emphasis is placed on instructional learning where children are provided with opportunities to engage in schoolwork and gain insights into their own performance based on teacher feedback (Denervaud, Knebel, et al., 2020). As this instructional learning takes hold, children learn to value correct answers and avoid errors (Denervaud, Knebel, et al., 2020). In contrast, the kindergarten environment introduces learning through more play-initiated activities (Morrison et al., 1997). While free play orientation may benefit children in many ways, it likely does not encourage the identification of errors on academic tasks as effectively as formal schooling (Denervaud, Knebel, et al., 2020).

Third, to determine whether the larger activation difference in performance monitoring in the P1 children could predict academic performance, we investigated the longitudinal coupling between these variables. We found borderline significant positive correlations between the change in activation in the left frontal cluster with performance on the math pack. This is in line with Kim et al. (2016) who found that stronger math skills (as well as reading skills) predicted stronger ERP component related to response monitoring. Further support for our finding stem from previous adult EEG research that found a larger ERN was significantly correlated with better academic performance as measured by student transcripts (Hirsh & Inzlicht, 2010). Given that monitoring one’s own performance is a key aspect of self-regulation, the authors interpreted that individuals with a greater ability to monitor engage in self-regulatory behaviours that are important for academic success (Pintrich & De Groot, 1990). It is however important to note that the change-change association between the left frontal cluster activation and math performance did not survive the formal model comparison. Therefore, the result needs to be interpreted with caution and stands for replication test. Future studies need to be better powered in terms of sample size. Hertzog et al., (2006) evaluated the statistical power of latent change score models and found even with large sample sizes and multiple measurement occasions, statistical power to detect covariance in change remains low. Given the modest sample size of the present study coupled with the inclusion of only two measurement occasions, we likely did not have sufficient statistical power to detect meaningful relationships, even when present.

Finally, we found that for KG children only, those who began the study with better performance on the CDT task showed a greater increase in response monitoring activation across the year. We did not predicted this result but it seems interesting, given that the KG children, at the mean level, did not show a significant change in activation difference across time. One interpretation for this finding relates to the interplay between children’s individual characteristics and the schooling / kindergarten environment. We posit that the schooling environment may have facilitated all school children, regardless of their starting point, to become more sensitive to task accuracy and error, leading to a mean change in brain activation across the year associated with stronger response monitoring. On the other hand, for the reasons highlighted above, kindergarten children may encounter less explicit instruction. Only those who are already advanced at the start, presumably by eliciting more advanced interaction with adult caregivers, show a change in brain activation associated with more efficient response monitoring. Future studies should test this postulation by getting more direct measurement of social/instructional environment of children.

To conclude, the present study is the first to use a cut-off design to assess the impact of one year of schooling on both response inhibition and response monitoring, and relate these differences to measures of academic achievement. No significant differences in response inhibition were found between the two groups of children. However, for response monitoring, after one year of schooling P1 children showed a greater activation difference than KG children. Functionally, this activation difference was associated with better performance on the go/no-go task. When relating to broader measures of academic achievement, we found only borderline significant associations between response monitoring and mathematical ability, preliminarily suggesting some functional relevance for school performance. Taken together, our findings highlight the role of the school environment in shaping the development of the neural network underlying monitoring of one’s error.

## Data Availability

All group-level data and analysis scripts have been made publicly available via the Open Science Framework and can be accessed at https://osf.io/nyjqe/. Further information regarding the fNIRS pre-processing and analysis are available upon request.

## Ethical Approval Statement

This research was approved by the General University Ethics Panel (GUEP 375a) at the University of Stirling.

## Acknowledgements

We would like to thank all the children and their parents who participated in the SAND project, as well as the schools and nurseries who distributed our study information.

## Footnotes

1 This project was pre-registered on As.Predicted.org (#34866). We initially planned to also examine the relationship between performance/neural activation of the go/no-go task with another behavioural EF task that taps into cognitive flexibility (heartsand-flowers task). However, due to an error in task programming, the data from the hearts-and-flowers task could not be interpreted. Therefore, we focused on children’s performance and brain response on the go/no-go task, and relate these to measures of academic achievement.

