## Supplementary Material for "Disentangling Age and Schooling Effects on Inhibitory Control Development: An fNIRS Investigation"

**Supplementary Materials**

*Academic Achievement in KG children*

*Right/left frontal cluster and academic achievement.* Bivariate longitudinal models were fitted for the response monitoring activation in the right frontal (or left frontal, respectively) and (1) vocabulary scores (2) math pack and (3) phonemes pack. The difference in activation in the left frontal cluster at T1 was positively correlated with vocabulary scores at T1. To follow up on this finding, the baseline correlation was constrained to be 0 which led to a significant drop in model fit $\Delta x^2 = 10.953$, $\Delta df = 1$, $p = .001$. Vocabulary scores at T1 also negatively predicted the change in activation in the left frontal cluster, suggesting KG children with better vocabulary performance at T1 showed less change in activation over time. To follow up on this finding, the coupling pathway was constrained to be 0 which led to a significant drop in model fit $\Delta x^2 = 6.119$, $\Delta df = 1$, $p = .013$. Parameter estimates are shown in Supplementary Table 1. The difference in activation at T1 positively predicted phoneme pack scores at T2 ($p$=.002), suggesting KG children who showed a greater difference in activation at T1 also showed better performance on the phonemes pack. To follow up on this finding, the pathway was constrained to be 0 which led to a significant drop in model fit $\Delta x^2 = 8.498$, $\Delta df = 1$, $p = .004$. No other cross-domain parameters were found to be significant in these models.

**Supplementary Table 1.** Bivariate couplings between left frontal activation and vocabulary for KG children.

| Bivariate Couplings | |
|---|---|
| Intercept covariance $\rho_{X1X2}$ | 65* (23.07) |
| Vocabulary onto left frontal change $Y1_{X2\Delta X1}$ | -1* (.37) |
| Left frontal change onto vocabulary $Y2_{X1\Delta X2}$ | -.04 (.03) |
| Change-change covariance $\rho_{\Delta X1\Delta X2}$ | 2.61 (20.48) |

Standard errors are in parentheses.

* Asterisks denote significance at $p$ <.05 level.
